# A probabilistic gene expression barcode for annotation of cell-types from single cell RNA-seq data

**DOI:** 10.1101/2020.01.05.895441

**Authors:** Isabella N. Grabski, Rafael A. Irizarry

## Abstract

Single-cell RNA sequencing (scRNA-seq) quantifies gene expression for individual cells in a sample, which allows distinct cell-type populations to be identified and characterized. An important step in many scRNA-seq analysis pipelines is the annotation of cells into known cell-types. While this can be achieved using experimental techniques, such as fluorescence-activated cell sorting, these approaches are impractical for large numbers of cells. This motivates the development of data-driven cell-type annotation methods. We find limitations with current approaches due to the reliance on known marker genes or from overfitting because of systematic differences between studies or batch effects. Here, we present a statistical approach that leverages public datasets to combine information across thousands of genes, uses a latent variable model to define cell-type-specific barcodes and account for batch effect variation, and probabilistically annotates cell-type identity. The barcoding approach also provides a new way to discover marker genes. Using a range of datasets, including those generated to represent imperfect real-world reference data, we demonstrate that our approach substantially outperforms current reference-based methods, in particular when predicting across studies. Our approach also demonstrates that current approaches based on unsupervised clustering lead to false discoveries related to novel cell-types.

## 1 Introduction

Single-cell RNA sequencing (scRNA-seq) quantifies gene expression at the level of individual cells, rather than measuring the aggregated gene expression in a biological sample containing millions of cells, as is done with bulk RNA-sequencing. This improved granularity permits the identification or discovery of distinct populations of cell-types within the tissues under study. To effectively accomplish this, it is important to annotate cells reliably by known cell-types, especially cells that are present in many tissues, such as immune system cells. Fluorescence-activated cell sorting (FACS) can be used prior to the sequencing step to physically sort cells from a mixed sample into their cell-type populations. While generally regarded as highly accurate, FACS-sorting has limited throughput, and thus is impractical when sequencing large numbers of cells. As a result, there is a need for data-driven approaches to annotate cell-types.

Current methods fall into one of two categories, which we will refer to as *clustering-based* and *reference-based*. In clustering-based methods, the more widely used approach, the target cells are first grouped using an unsupervised clustering algorithm (for example, [1, 2, 3, 4]). Next, differential expression analysis is used to identify genes that are uniquely expressed in each group and compared to known cell-type-specific marker genes to annotate the group as a particular cell-type. Although any clustering algorithm can be used in this type of method, we focus primarily on Seurat’s clustering algorithm [4], since it is popularly used in this way for cell-type identification.

Reference-based methods (for example, [5, 6, 7, 8, 9]) use supervised learning approaches in which the target cells are compared to reliably annotated, such as by FACS-sorting, reference data for each cell-type of interest, and each target cell is annotated using the *closest match*. Approaches to defining *closest match* vary. Many of these supervised methods are based on complex and hard-to-troubleshoot machine learning algorithms, such as Xgboost [7] and deep neural networks [10]. As a result, unexpected systematic differences between training and test sets may lead to over-fitting. In addition, some of the most popular methods rely on marker genes to guide the determination of the closest match [6, 5]. However, reliable marker genes are not always known for every cell-type of interest.

In this work, to avoid reliance on marker genes and minimize over-fitting, we consider all genes as potentially informative and develop a latent variable model that characterizes cell-types by the probabilities of genes being in expressed or not-expressed states, a probabilistic barcode. Other sources of within-state variability, such as that introduced by batch effects, are modeled by conditional distributions. Exploratory data analysis, described in the Results Section, demonstrates the need for these to be gene-specific distributions. We therefore implement a two-stage procedure: first, we estimate gene-specific parameters using a fixed public database, and second, we estimate cell-type-specific probabilities, the barcodes, for each cell-type using training data. To classify cells into cell-types, we fit this model and use the resulting fit to compute posterior probabilities.

Here, we describe the datasets used to build and assess our method, provide a detailed description of our approach, demonstrate its advantages, and show the limitations of existing approaches. Our model-based approach also helps bring to light the false positive results produced by current approaches to discovering new cell-types, by examining results from the Allen Brain Map [11].

## 2 Results

### 2.1 Datasets

#### 2.1.1 PanglaoDB database

To motivate and fit gene-specific distributions across tissues and cell-types, we used the PanglaoDB database [12], which provides publicly available scRNA-seq data from a diverse set of experiments. We considered only the datasets corresponding to non-tumor samples from humans. This yielded 218 datasets comprising a total of 3,389,679 cells, with each dataset representing one cell-type or tissue-type.

#### 2.1.2 Cell atlas data

To study an example of computationally derived cell clusters in a popular atlas, we considered human brain data from the primary motor cortex. This data was clustered by the original study using `scrattch.hicat` [11].

#### 2.1.3 Assessment data

To benchmark our approach against existing methods, we constructed three assessment datasets. We selected datasets for which we were highly confident of the accuracy of assigned cell-type labels. For each of these, we constructed a *main* dataset from one or more published studies. This dataset was split into a training set and a test set. A second test set was formed using data from a separate study not included in the main dataset. We refer to these two test sets as the *withheld test set* and the *external test set*, respectively. Note that over-training due to study-specific biases and batch effects will result in better performance in the withheld test set compared to the external test set.

To mimic situations in which finding large and accurately labeled training datasets is challenging, we constructed four versions of the training set. The *complete sample* version included the entire training dataset. The *small sample* version was constructed by randomly selecting 50 cells from each reference cell-type from the complete sample. To construct the *contaminated* version, 25% of reference cells were assigned a label at random. Finally, to construct the *downsampled* version, we downsampled a subset of the reads from the complete sample to 50% coverage. Specifically, we downsampled all cells corresponding to a cell-type present in the external test set.

##### PMBC dataset

The first main dataset combined four different FACS-sorted human peripheral blood mononuclear cell (PBMC) datasets from 10X Genomics. In particular, we combined 11,113 CD4 cells, 10,109 CD8 cells, 2,512 CD14 cells, and 8,285 NK cells [13]. To construct the external test set, we combined 425 isolated CD14 cells each from the blood of two different donors from a different study [14], to result in a total of 850 test cells.

##### Colon dataset

The second main dataset combines three different human colon datasets [15, 16, 17]. The first provided 4,472 experimentally purified colon epithelial cells [15]. Although these were derived from patients with intestinal tumors, the tissue was collected at least 10 centimeters away from the tumor border. The second provided 21,248 healthy mesenchymal stromal cells [16] from the colon, purified through MACS-depletion. The third provided 3,461 FACS-sorted B cells from colon samples [17] from pediatric patients with inflammatory bowel disease. To construct the external test set, we used 1,000 each of epithelial, stromal, and B cells from healthy colons from a different study [18]. This study assigned cell-type labels using a combination of clustering and expert evaluation of differentially expressed genes. We therefore only considered epithelial, stromal, and B cell-types, as they are sufficiently distinct from one another to result in accurate labeling. Although the study identified more specific sub-types within the epithelial and stromal cells, we just used the high-level labels “epithelial” and “stromal” for such cells, instead of the finer-grained sub-type labels, to ensure accuracy.

##### Brain dataset

For the third dataset, we used a subset of the non-neuronal cells from the previously described cell atlas data. Specifically, we used 2,942 oligodendrocytes, 568 astrocytes, and 108 microglia/PVM cells. These cell-type labels are computationally derived by the original study. Due to the small number of microglia cells available, we only used oligodendrocyte and astrocyte cells to construct the withheld test set. For the external test set, we used an experimentally purified dataset [19] consisting of microglia.

All our datasets were processed with 10X Genomics technology using unique molecular identifiers (UMI).

### 2.2 Clustering-based approaches artificially identify new cell-types

We used the PBMCs dataset to investigate the relationship between the number of clusters and the dataset size when clustering with Seurat [4]. In particular, we applied the Louvain clustering algorithm [20] implemented in Seurat v3 to successively subsetted versions of the PBMC dataset, ranging from a total of 80 cells (20 per cell-type) to a total of 2,000 cells (500 per cell-type). Although there are four cell-types represented in this dataset, the number of clusters found ranges from three to six, with a general increasing trend with dataset size (Figure 1(A)). Only a narrow window of dataset size (roughly between 500 and 1,000 total cells) corresponds to the correct number of cell-types.

**Figure 1:**
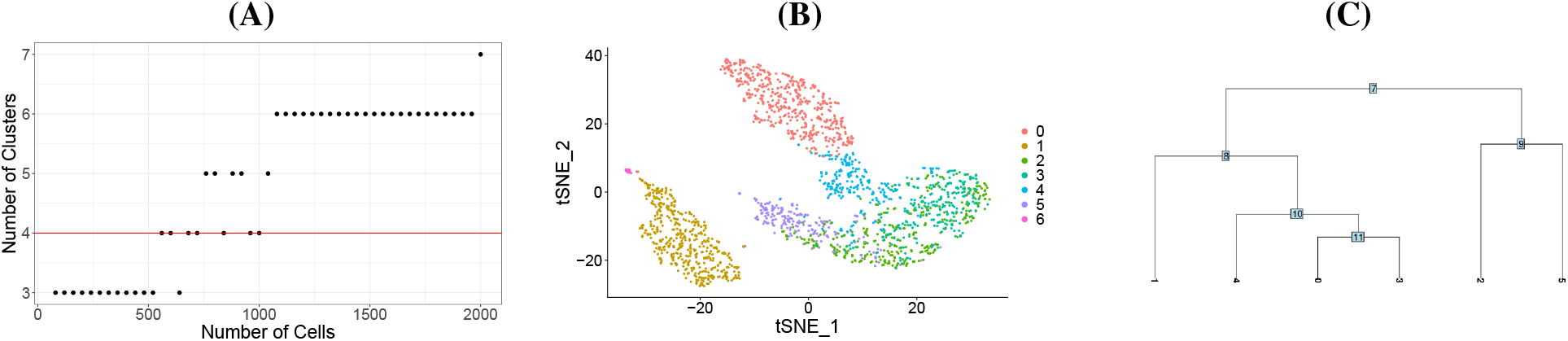
Applying Seurat clustering to successive subsets containing equal amounts of each of the same four cell types (CD4 T-cells, CD14 T-cells, CD8 T-cells, and NK cells) from the PBMC reference data incorrectly identifies more cell-types with increasing dataset size. (A) Number of clusters identified by Seurat plotted against the number of cells included in the analysis. The red line denotes the true number of cell-types. (B) tSNE plot of the six clusters, denoted with different colors, identified in the largest dataset size, which consisted of 2,000 cells. (C) Dendrogram of the six clusters identified in the largest dataset size, constructed using the average cell of each cluster. The leaf labels denote the cluster numbers.

We further examined the six clusters found when applying the algorithm to the 2,000 cells using t-distributed stochastic neighbor embedding (tSNE), which did not show separation among the six identified clusters (Figure 1(B)). Furthermore, the clusters are hierarchically related in ways that do not correlate with the true cell-type identities of the cells within those clusters (Figure 1(C)). For example, clusters 2 and 5 represent a division from a larger group, but in fact, nearly every cell in both of those clusters is a CD4 cell. Note that clusters 0, 3, and 4 are all close, but the cells in cluster 0 are denoted as nearly half CD8 and half CD14 by FACS-sorting, with almost all in the cells in cluster 3 corresponding to CD8 cells and almost all the cells in cluster 4 corresponding to CD14 cells. Hence, these extra divisions cannot be simply interpreted as the result of identifying sub-types within larger categories of cell-types. Only cluster 1, which consists almost entirely of NK cells, corresponds to a single cell-type with specificity. These findings suggest that the identity assigned to a given cell depends on the size of the dataset, in a way that is not representative of the actual cell-types in the data.

#### 2.2.1 Marker genes are unreliable due to sparsity

Clustering-based methods rely on marker genes. We therefore examined the reliability of such marker genes for labeling in scRNA-seq data. To do so, we used a subset of the reference PBMC dataset as an example and looked at the counts for externally validated marker genes for each cell type. Specifically, we selected marker genes for CD4 (IL7R, CD4, CTLA4, FOXP3, IL2RA, PTPRC), CD14 (CD14, LYZ, FCGR3A, CD68, S100A12), CD8 (CD8A, CRTAM, NCR3, CD3D), and NK cells (GNLY, NKG7, PRR5L, S1PR5, NCAM1) that have been curated and well-established in the literature as discriminating among cell types [21, 22, 4]. This list of marker genes is representative of those that might be chosen by a user applying a marker-based method to a PBMC dataset.

Among these genes, GNLY and NKG7 are highly sensitive and highly specific to NK cells, as is LYZ for CD14 cells (Figure 2). However, we observe 0 cells with non-zero counts even for marker genes that are known to be expressed (Figure 2). Specifically, both CD14 and CD4 cells had markers with non-zero counts in 0% of cells, and the remaining two cell-types, NK and CD8, both had markers with non-zero counts in as low as 4% of cells. Among CD4 cells in particular, the highest proportion of non-zero counts for any of its markers was only 38% (Figure 2). In summary, we found that most marker genes are not definitively and specifically present in scRNA-seq data.

**Figure 2:**
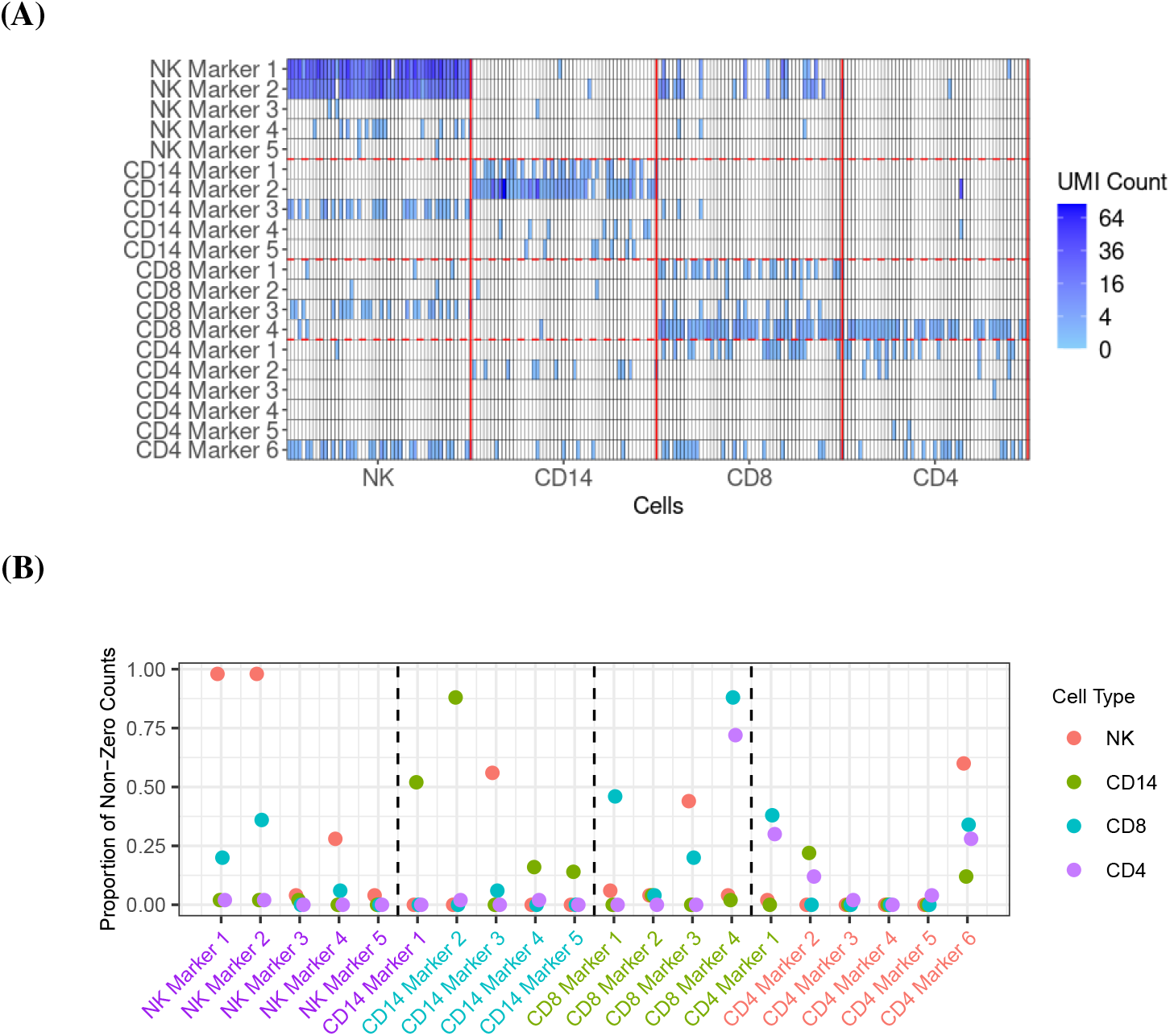
PBMC data from 20 canonical marker genes show that they are unlikely to be observed. (A) UMI counts, shown in shades of blue, for marker genes in 50 randomly selected cells from each pertinent cell-type. (B) Proportion of non-zero UMI counts for the same canonical marker genes and each cell-type, which is denoted with color.

### 2.3 Model-based probabilistic classifications provide substantial improvements

Given the limitations of clustering-based approaches, we considered reference based approaches instead. We denoted the unknown cell-type for cell *i* as *Y*_*i*_ and the observed gene counts as **x**_*i*_ = *x*_*i*1_, …, *x*_*iJ*_ with *J* the total number of genes. Supervised, or reference-based, approaches are generally based on using training data to estimate the conditional probability of *Y*_*i*_ given gene counts **X**_*i*_ = **x**_*i*_. Specifically, for each cell *i*, we want to estimate the conditional probability

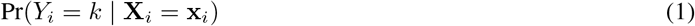

and then select the *k* that maximizes it. There are several proposed approaches to estimating this multivariate conditional probability [23]. However, the large number of genes and the sparsity of the data, demonstrated above in Section 2.2.1, make this challenging. Furthermore, batch effects introduce systematic biases that make gene counts difficult to compare across studies [24]. Here, we developed an approach that tackles these challenges by 1) using a conditional Poisson model to account for the sparsity and differences in coverage, 2) introducing a parsimonious parametric model that assumes counts are independent across genes once we condition on cell-type, and 3) providing robustness to batch effects by assuming a latent state model for each gene that models within-state variability in a way that down-weights its influence on prediction.

We start by using Bayes rule to rewrite the posterior (1) as:

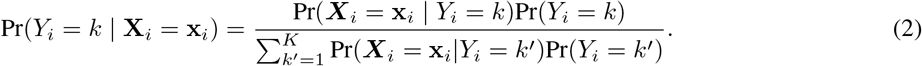

We then assumed that each of the *K* possible cell-types is equally likely: Pr(*Y*_*i*_ = *k*) = 1*/K*. To make the model parsimonious, we assumed that, conditional on the cell-type, the *X*_*ij*_ are independent across genes *j*:

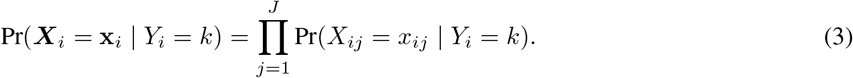

To account for sparsity, we assumed that for any cell-type *k, X*_*ij*_ can be modeled as

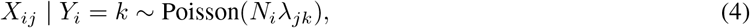

with *λ*_*jk*_ proportional to the expected gene expression for gene *j* in cell-type *k* and 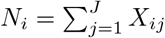 the total observed counts across all genes in cell *i*. A challenge with supervised approaches is that to train the model, we have to estimate *λ*_*jk*_ from data in the training set, which suffers from the limitations listed above: 1) we often have few cells available for estimation, 2) data can be sparse, and 3) batch effects result in systematic biased estimates that result in unsuitable across-study performance. We developed an approach that tackled these limitations by 1) assuming that the biological information is represented by expressed (on) and not-expressed (off) latent states and 2) accounting for other sources of variability with a random variable *λ*_*j*_ that follows a parametric distribution and assuming the *λ*_*jk*_ were realizations of this random variable. Next, we describe how we developed this parametric model.

#### 2.3.1 Count distributions are bimodal and gene-specific

To determine a parametric form for the gene-specific distributions of *λ*_*j*_, we leveraged the fact that the PanglaoDB database offers large numbers of cells for each cell-type *k*, and defined estimates

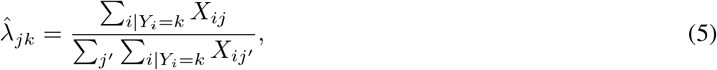

assumed that 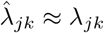, and used exploratory data analysis. Specifically, we used 3,389,679 cells from 218 cell-types and tissues from the PanglaoDB database to obtain precise estimates 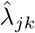 for *k* = 1, …, 218. We found that many genes showed a clear bimodal distribution for these rates when examined across cell-types, consistent with our latent state model assumptions (Figure 3). Furthermore, we noted that the centers of the off and on distributions vary by gene, consistent with results previously observed using gene expression data from the Gene Expression Omnibus and Array Express [25]. Note that this implies that observed gene count rates are not comparable across genes. For example, note that some values associated with the off distribution for RPL31P7 would be categorized as on for genes such as AC140076.1 (Figure 3).

**Figure 3:**
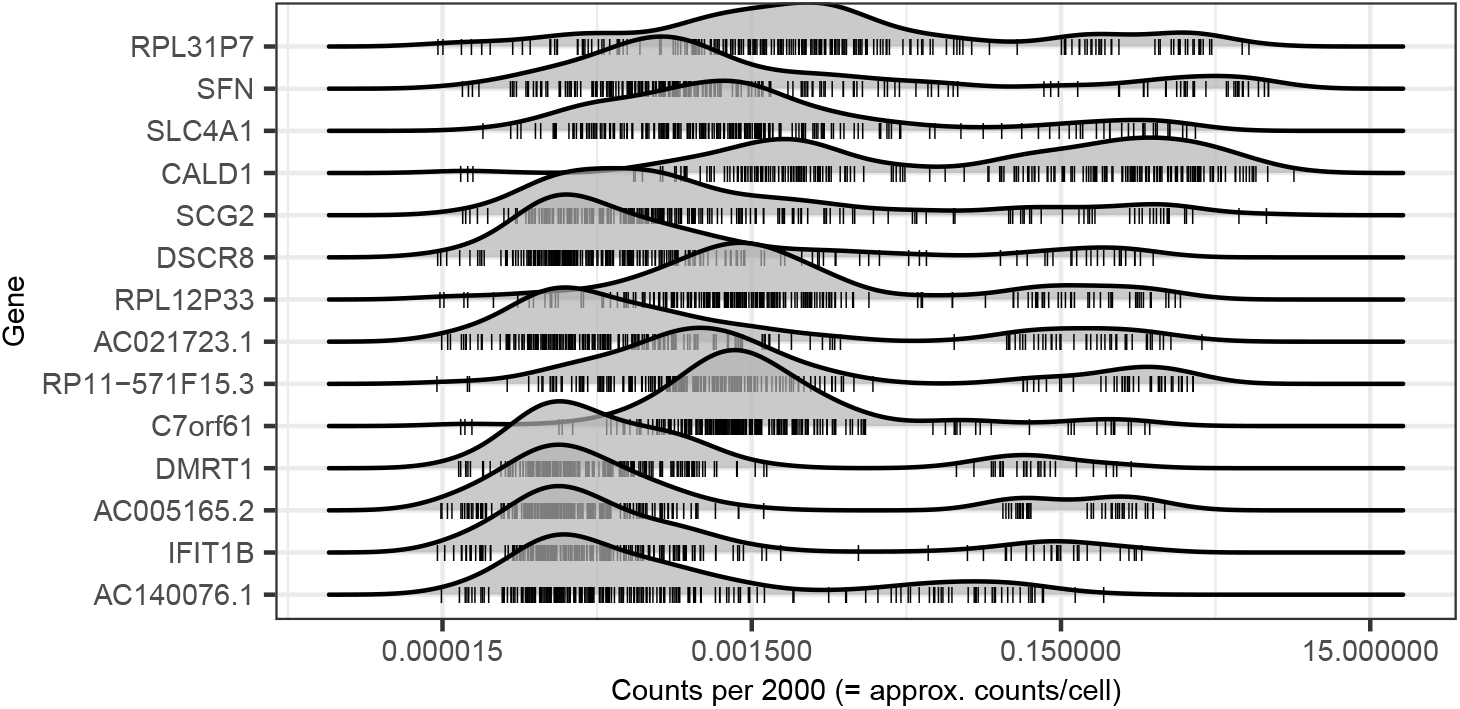
Example of density plots of genes with bimodal expression distributions across cell- and tissue-types. Every tick mark represents the rate of that gene in a particular cell- or tissue-type. The centers of the on and off distributions can be seen to vary by gene.

Further data exploration (Supplementary Figure 1) led us to assume the off distribution was best modeled by a mixture of exponential and log-normal distributions, with the log-normal component accounting for low counts consistent with a non-zero background level of expression distinctly lower than the expressed state, and the exponential accounting for counts that were mostly zeros, consistent with practically no expression (Supplementary Figure 2). We therefore further divided the off state into two. We represented the latent structure by introducing the unobserved variable *Z*_*jk*_ that can be one of three states, *off-low, off-high*, or *on*, for each gene *j* in each cell type *k*. For simplicity of exposition, we will refer to the *off-high* state as simply *off*. Note that if the *Z*_*jk*_s were observed, the vector ***Z***_*k*_ can be considered a gene expression barcode that uniquely identifies cell-type *k*. The main challenge of our approach is estimating these *Z*_*jk*_. We connect the unobserved *Z*_*jk*_s to the observed ***X*** by modeling the distribution of *λ*_*j*_ as

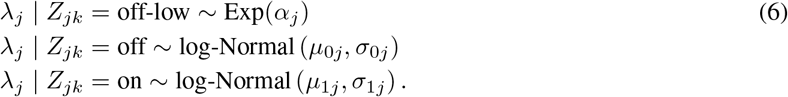

Here, *µ*_0*j*_ and *µ*_1*j*_ represent gene-specific means of the off and on distribution respectively. The gene-specific standard deviations *σ*_0*j*_ and *σ*_1*j*_ quantify the variability that accounts for the fact that we observe different rates for tissues in which gene *j* is off and tissues in which gene *j* is on, respectively. This includes variability introduced by batch effects.

As described in detail in the Methods Section, our approach to estimating this model is to first estimate the gene-specific parameters *α*_*j*_, *µ*_0*j*_, *µ*_1*j*_, *σ*_0*j*_, and *σ*_1*j*_ using the PanglaoDB database. These parameter estimates are then *frozen* and used by our algorithm when training on new datasets. With these parameter estimates in place, we can then obtain estimates 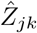 by applying the model (6) to the training data and then using these to estimate the posterior probability (1) for the test set and subsequently classify each cell into a cell-type. The vector 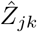 can be considered a probabilistic barcode for each cell-type *k*.

### 2.4 Our approach improves current reference-based methods

We compared our method to six leading classification methods: [9], scmap [8], CaSTLe [7], SingleR [26], Garnett [6], and CellAssign [5]. For each of these methods, we processed the reference and test datasets as recommended in their respective documentations, and for the methods requiring markers (Garnett and CellAssign), we identified six markers for each cell-type using `scran` [27] on the reference dataset. This was done for consistency because not every cell-type under consideration has readily available, independently verified marker genes. For the runs with Garnett, we used Garnett’s built-in marker diagnostic function to assess the marker genes on the corresponding reference data, and we dropped any high ambiguity markers. The one exception was for the Brain dataset with the contaminated reference, where all top markers for microglia were flagged for high ambiguity and so we retained all of them.

The results of the benchmark are shown in Table 1. Our approach performed well universally: the lowest accuracy achieved in the external validation, across all comparisons, was 0.982. No other method achieved this. CHETAH, scmap, and CaSTLe had strong performance on some withheld sets but much weaker performance on the external sets, a consistent outcome for complex machine learning methods susceptible to over-training. Moreover, methods such as Garnett and CellAssign performed poorly on the Brain dataset, which was the most challenging dataset given the similarity of cell-types and low numbers of available reference cells. Because Garnett and CellAssign primarily use marker genes to guide their classifications, they may be especially sensitive to such data since any distortions to the marker gene counts could have a large effect on the final prediction.

**Table 1:** Classification accuracy for our approach, the probabilistic barcode, and six leading methods, as evaluated on three main datasets (PBMC, Colon, and Brain) with two types of test sets (withheld and external) each. Four types of reference data were used, indicated in the third row as complete sample (CS), downsampled (D), small sample (S), and contaminated (C). The last column shows the worst accuracy obtained on any external dataset for each method.

|  | PBMCs |  |  |  |  | Colon |  |  |  |  | Brain |  |  |  |  | Worst |
| --- | --- | --- | --- | --- | --- | --- | --- | --- | --- | --- | --- | --- | --- | --- | --- | --- |
|  | Withheld | External |  |  |  | Withheld | External |  |  |  | Withheld | External |  |  |  |  |
|  | CS | CS | D | S | C | CS | CS | D | S | C | CS | CS | D | S | C |  |
| Barcode | 0.975 | 0.985 | 0.985 | 0.982 | 0.984 | 0.990 | **0.998** | **0.998** | **0.998** | **0.995** | **1.000** | **1.000** | **1.000** | **1.000** | **1.000** | **0.982** |
| CHETAH | 0.943 | 0.911 | 0.982 | 0.981 | 0.991 | 0.910 | 0.425 | 0.541 | 0.432 | 0.740 | 0.970 | 0.945 | 0.903 | 0.860 | 0.000 | 0.000 |
| scmap | 0.700 | 0.769 | 0.002 | 0.407 | 0.379 | 0.833 | 0.937 | 0.769 | 0.908 | 0.574 | 0.965 | 0.001 | 0.000 | 0.510 | 0.001 | 0.000 |
| CaSTLe | **0.985** | 0.986 | 0.948 | 0.981 | 0.981 | 0.997 | 0.710 | 0.804 | 0.743 | 0.612 | **1.000** | 0.607 | 0.999 | 0.658 | 0.269 | 0.269 |
| SingleR | 0.945 | 0.955 | 0.878 | 0.972 | 0.875 | **0.983** | 0.841 | 0.848 | 0.954 | 0.824 | **1.000** | 0.962 | 0.945 | 0.942 | 0.942 | 0.824 |
| Garnett | **0.880** | **1.000** | **1.000** | **1.000** | **1.000** | 0.837 | 0.938 | 0.947 | 0.563 | 0.819 | 0.710 | 0.861 | **1.000** | 0.000 | **1.000** | 0.000 |
| CellAssign | 0.738 | 0.972 | 0.979 | 0.982 | 0.966 | 0.973 | 0.982 | 0.982 | 0.984 | 0.986 | **1.000** | 0.000 | 0.000 | 0.000 | 0.000 | 0.000 |

In cases where the reference data were modified, the effect on performance was quite variable across methods and datasets. Nevertheless, each method besides ours displayed worsened performance in at least some of the assessments.

For example, for scmap, this occurred the most with downsampling, whereas for CaSTLe, this happened under label contamination. The most consistently robust method, aside from our approach, was SingleR, which never achieved lower than 82.4% accuracy.

#### 2.4.1 Overclustering may lead to artificially identified cell-types in the Allen Brain Map

The Allen Brain Map [11] used clustering to identify 127 distinct cell-types at the lowest level of the hierarchy, of which 44 correspond to excitatory neuron sub-types and 72 correspond to inhibitory neuron sub-types. To explore the possibility that some of these findings were the result of over-clustering, we examined data from four of the 44 excitatory sub-types and four of the 72 inhibitory sub-types. Specifically, we 1) took a random sample of 50 cells from each of the cell-types Exc L5-6 FEZF2 SH2D1B, Exc L5 THEMIS LINC01116, Exc L5 THEMIS RGPD6, Exc L5-6 FEZF2 FILIP1L cell-types, Inh L5-6 SST PAWR, Inh L3-5 SST GGTLC3, Inh L3-5 SST OR5AH1P, and Inh L5 SST RPL35AP11, 2) obtained gene expression counts for each cell of 100 genes identified as marker genes for all cell-types by the Allen Brain Map project, and 3) computed a Poisson-based dissimilarity measure [28] over these genes for each pair of these 400 cells. Note that the 100 genes were chosen as those with the largest summed counts across the 8 clusters out of the 206 total genes indicated as markers in the original study. A clear difference in gene expression patterns was observed between excitatory and inhibitory neurons, but we saw no clear differences between the new cell-types within each of these two (Figure 4A). Furthermore, the average expression profile of each of these cell-types, taken over all cells with these labels in the database, does not show clear differences between the new sub-cell-types (Figure 4B).

**Figure 4:**
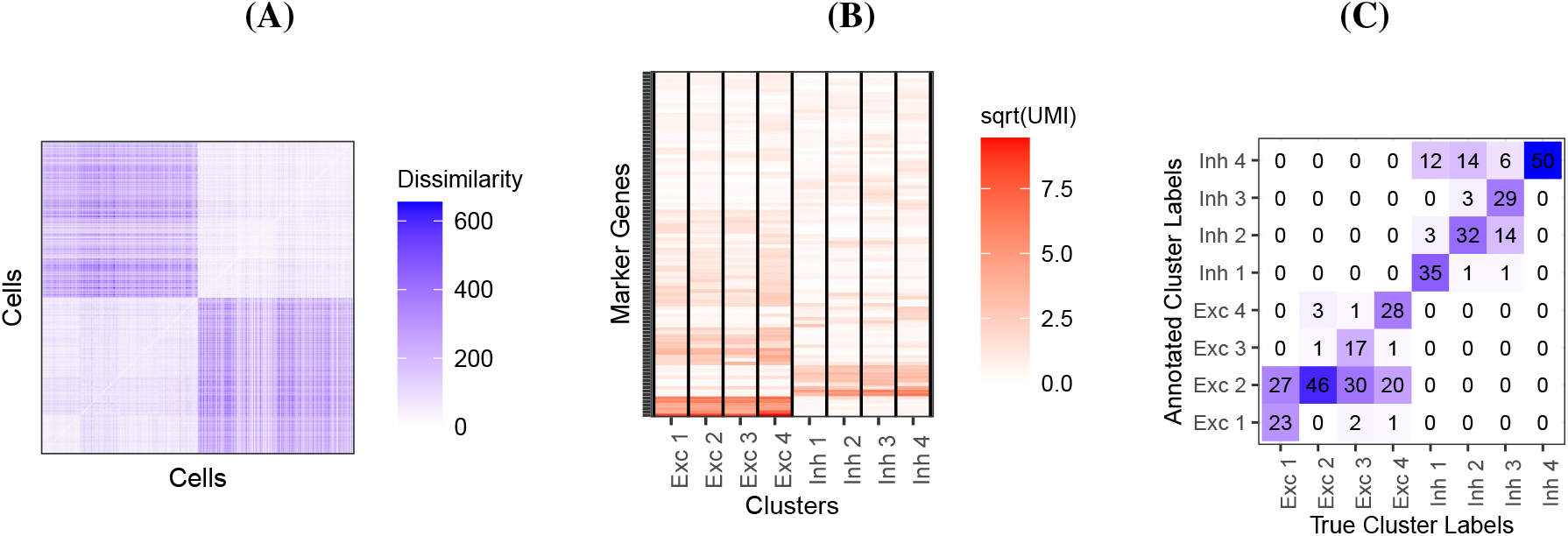
Evidence of artificially identified cell-types in the Allen Brain Map. For ease of visualization, we label sub-types by “Exc” or “Inh,” followed by their ranking among themselves in number of assigned cells. (A) Dissimilarities between random samples of cells belonging to four excitatory neuron cell-types and four inhibitory neuron cell-types in the Brain dataset, defined by clustering in the original study. Values are computed using 100 of the 206 total marker genes observed over all cell-types in the dataset. A larger value corresponds to more dissimilarity. (B) Square root of average UMIs observed in each cell of the same random samples from each cluster for the same set of 100 marker genes. (C) Contingency table showing the agreement between the annotated cluster labels under our approach and the study’s true cluster labels, for the same random samples. The model was trained using the cells not selected as part of the random samples.

We additionally explored these sub-types under our approach, by training our model on the cells not included in the random samples and then classifying these hold-outs. We found that our method perfectly distinguishes between excitatory and inhibitory neurons, but many cells were not assigned their specific sub-type label (Figure 4C). Instead, 61.5% excitatory neurons were assigned to Exc L5 THEMIS RGPD6, and 41% inhibitory neurons were assigned to Inh L5-6 SST PAWR. This result was consistent with there being no novel sub-types among this set, and Exc L5 THEMIS RGPD6 and Inh L5-6 SST PAWR representing the excitatory and inhibitory neurons, respectively.

#### 2.5 Gene-specific model predicts marker gene effectiveness

Fitting model (4) permitted us to explain why some markers are more effective than others, and to identify new marker genes. For example, note that for the FOXP3 gene, a marker we found to be reasonably effective for CD4 cells, the on distribution is clearly distinguishable from the off distribution (Figure 5). In contrast, for the gene named CD4, which we found to be an ineffective marker despite having a larger rate than FOXP3 among CD4 cells in our PBMC dataset, we see less clear separation of the on and off distributions. Furthermore, markers such as PTPRC and IL7R have well-separated distributions but are on in many tissue types, which makes them less effective as well. Our method was also useful for defining new markers: for a given cell-type *k* ^*′*^ we search for genes *j* for which Pr(*Z*_*jk*_ = on) is high, but Pr(*Z*_*jk*_*′* = on) is low for most or all *k*^*t*^ = *k*. Using this approach, we identified genes NDUFA13, PPIB, PCBP1, and POLR2F as effective markers for CD4 cells (Figure 5), for example.

**Figure 5:**
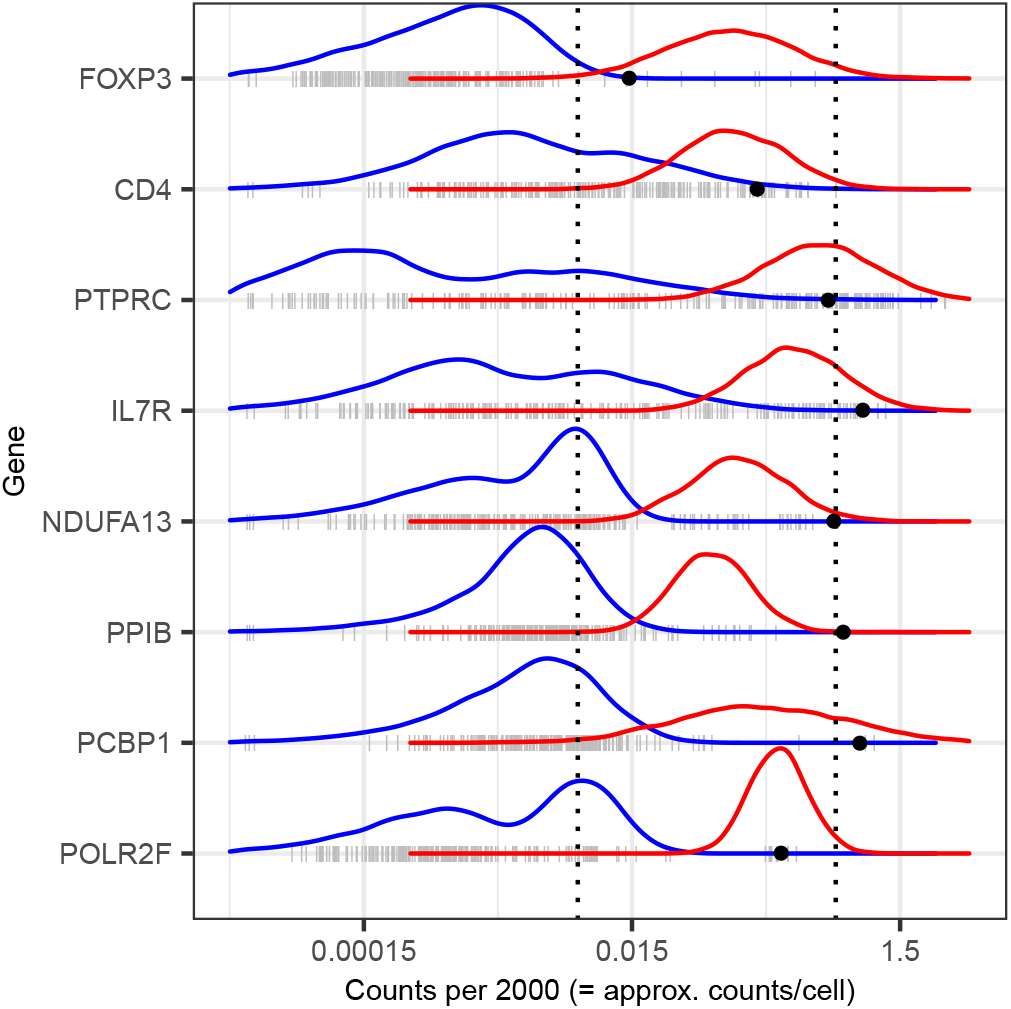
Distributions of the rates of four canonical CD4 markers (top four) and four CD4 markers we identified (bottom four) across cell- and tissue-types, against the fitted off (blue) and on (red) components for each gene. Gray ticks show the rates of that gene in a particular cell- or tissue-type. Black points indicate the rates in CD4 cells from the PBMCs dataset.

## 3 Discussion

Currently, cell-type identification in scRNA-seq datasets is done with either clustering-based or reference-based methods. We showed that clustering-based methods can artificially identify more cell-types as the dataset size increases. We also showed that overclustering may be present in widely-used resources such as the Allen Brain Map. Moreover, clustering-based methods can depend on arbitrary user decisions such as the choice of marker genes, which is a problem for cell-types that do not have well-established and widely-accepted markers. Different users examining the same clusters might draw two different conclusions about the cell-type identities. We further showed that marker genes in scRNA-seq data can have varying reliability, even for well-studied cell-types. This method of direct annotation by a user additionally implies that probabilistic classification is not possible; a given target cell is assigned to some particular cell-type without any information about the certainty of this assignment. Finally, with clustering approaches it is difficult to specify the desired granularity of the cell-types. Since clusters are identified in an unsupervised manner, there is no differentiation between, for instance, a use case where it is enough to identify cells as *T-cells* and a use case in which identification at the level of T-cell subtypes is needed.

Reference-based methods avoid the pitfalls of clustering-based methods. The supervised approach assures that the desired granularity can be controlled with the choice of reference data, and avoid artifacts such as the number of distinct cell-types identified growing as the dataset size increases. Nevertheless, we showed that many of the currently available algorithms are susceptible to over-training due to study-to-study variability or batch effects. In addition, most methods do not provide probabilistic classifications. We introduced an approach that provides a solution to these challenges. First, we leverage thousands of genes, rather than only a few markers, to make the identifications, which provides robustness to the challenges introduced by sparsity. Second, we directly account for coverage in our model, allowing us to make reliable classifications even with varying coverage between the reference and test data. Finally, to account for study-specific biases and batch effects we assume a latent variable model, model unwanted variability with gene-specific distributions across cell-types, and represent each reference cell-type with a unique probabilistic barcode.

We demonstrated the advantages of our approach by assessing perfomance on several real-world datasets, with reference sets of varying quality. In particular, our approach maintained strong performance when the reference data had a small sample size, were downsampled, or had partially contaminated labels, whereas other methods suffered from poor performance under some of these scenarios. Since high-quality reference data can be difficult to obtain, these results suggest that our approach remains useful even if only low-quality or somewhat unreliable data are available. Moreover, our approach was successful at generalizing to external test sets. It is possible that the poor performance of the two marker-based methods (Garnett and CellAssign) on some of the datasets could be attributed to the choice of markers, and that a richer analysis to guide marker selection could have yielded better results. Even if better performance might be possible, this would require extra steps beyond what is part of the method. By contrast, our approach does not require additional inputs other than the reference data.

## 4 Methods

### 4.1 Estimating Gene-Specific Parameters

We started by estimating the parameters in model (6) using data from the PanglaoDB database. Because several genes had few tissues in the on state, and some genes, the housekeeping genes for example, had few tissues in the off state, to improve power, we borrowed strength across genes by imposing a prior distribution. Data exploration implied that gene-specific on and off means were correlated across genes, so we used a bivariate normal distribution with correlation *ρ*:

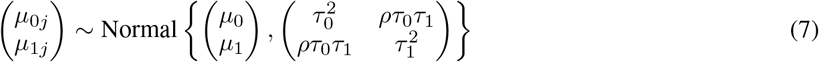

Here *µ*_0_ and *µ*_1_ represent the overall expected rate for genes that are off and on, respectively, and *τ*_0_ and *τ*_1_ quantify the variability in the gene-specific shifts.

In the next step, we used the 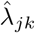 defined in (5) to estimate the parameters that define (6). We noted that the marginal distribution of the random variable *λ*_*j*_ can be written as

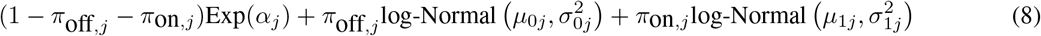

with *π*on_,*j*_ = Pr(*Z*_*j*_ = 1) and the same parameters as those that defined the distributions in (6).

We selected prior parameters *µ*_0_, *µ*_1_, *τ*_1_, *τ*_2_, and *ρ* using an empirical approach. Specifically, we considered housekeeping genes [29], for which the microarray-based gene expression barcode [25] estimated non-zero probabilities of being on across all healthy tissues. We then took the mean of the log-rates 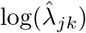, as defined by (5), for these genes in the PanglaoDB database as *µ*_1_, and the sample standard deviation as *τ*_1_. We next took all genes for which the microarray-based gene expression barcode estimated zero probabilities of being on across all healthy tissues and used expectation-maximization to fit a mixture of the off components, namely an exponential and a log-normal distribution, to all the rates 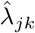 of this subset of genes in the PanglaoDB database. We used the mean of the log-rates for genes with 95% probability or higher of belonging to the log-normal distribution as *µ*_0_, and the sample standard deviation of these same genes as *τ*_0_. We set *ρ* = 0.5 based on exploratory data analysis.

With the prior distribution in place, we then fit the model to 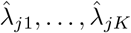 using the Expectation-Maximization (EM) algorithm for each gene *j*. To prevent label-switching and ensure interpretability of the model, we also set the constraints *µ*_0*j*_ *< µ*_1*j*_, *σ*_0*j*_ *>* 0.5, and *σ*_1*j*_ *>* 0.5 for all *j*. Details are in Supplementary Information Section 2. We considered the gene-specific parameters to be frozen going forward and denoted them with 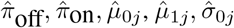 and 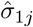.

### 4.2 Estimating the probabilistic barcode from training data

With estimates of the gene-specific distributions (6) in place, all we needed to produce a classification algorithm based on our estimate of the posterior probabilities (2) was an estimate of the *Z*_*jk*_ for each gene *j* and cell-type *k*.

For each cell-type *k*, we defined

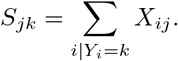

We can show that if (4) and (6) hold, then

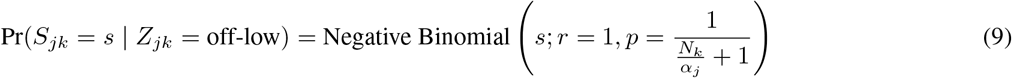

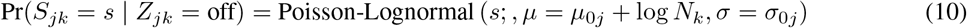

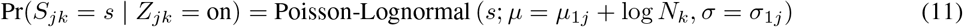

We can then plug-in our frozen estimates 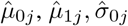, and 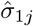 into 9 to define estimates of *Z*_*jk*_ as

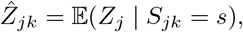

which we can determine by computing Pr(*Z*_*j*_ = *l* |*S*_*jk*_ = *s*) for each state *l* (off-low, off, on). These are found using Bayes rule:

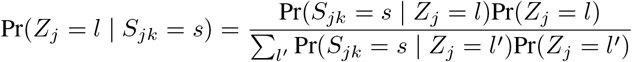

where we can plug-in our existing estimates 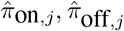, and 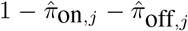 respectively for Pr(*Z*_*j*_ = *l*), and Pr(*S*_*jk*_ = *s* | *Z*_*j*_ = *l*) are computed as above.

The resulting probabilities constitute our cell-type-specific barcode. We can plug-in our frozen estimates and rewrite (6) as

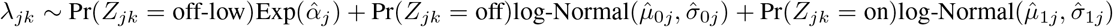

Then to classify each unknown cell, we first selected informative genes by considering only those exhibiting bimodal expression based on their gene-specific distributions. Specifically, we required a large enough difference between the off and on gene-specific means *µ*_1*j*_ −*µ*_0*j*_ *>* 1 and that the probability of gene being always off or always on was less than 5%: 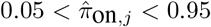. This filter yielded a set of 6,996 genes. Considering only these informative genes, we can evaluate the posterior probability of each test cell belonging to each cell-type of interest as in (2). Further details on implementation are in the Supplementary Methods and the code at https://github.com/igrabski/scRNAseq-cell-type.

