## Supplementary Information for "A probabilistic gene expression barcode for annotation of cell-types from single cell RNA-seq data"

December 22, 2020

### 1 Supplementary Figures

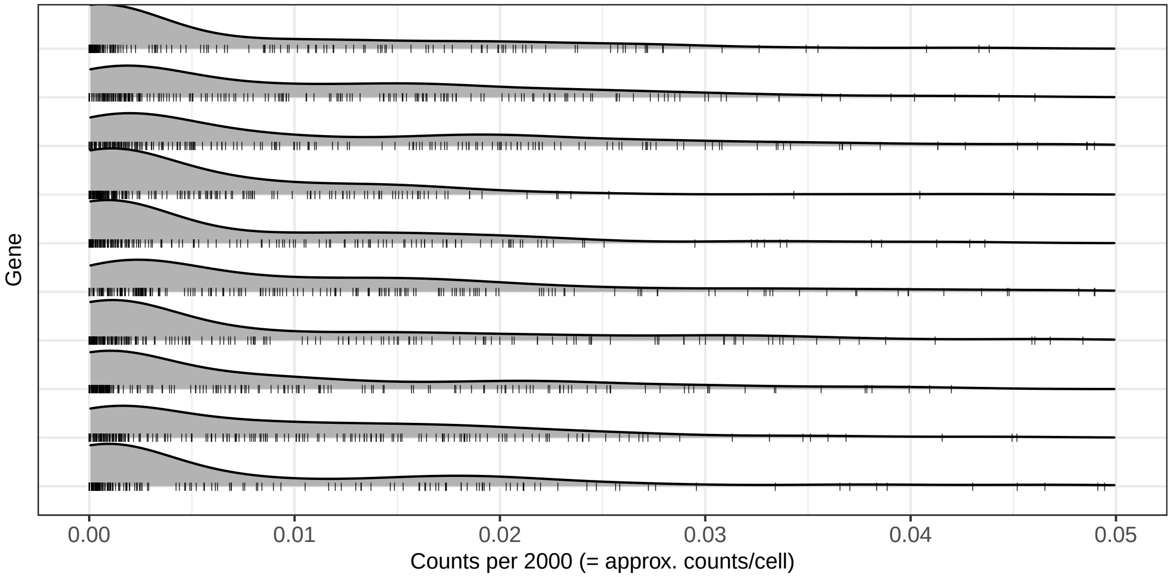


Figure 1: Examples of genes determined to be unexpressed based on the microarray barcode (i.e. probability of being on is 0 in every healthy tissue), demonstrating two off states: rates that are zero or near-zero, and rates that are low but not near-zero. These can be observed from the bimodal expression within these unexpressed genes.

### 2 Derivations for EM Algorithm

For computational simplicity, we define $\delta_{0j}, \delta_{1j}$ such that

$$\mu_{0,j} = \mu_0 + \delta_{0j}$$

and

$$\mu_{1,j} = \mu_1 + \delta_{1j},$$

where

$$\begin{pmatrix} \delta_{0j} \\ \delta_{1j} \end{pmatrix} \sim \text{Normal} \left\{ \begin{pmatrix} 0 \\ 0 \end{pmatrix}, \begin{pmatrix} \tau_0^2 & \rho\tau_0\tau_1 \\ \rho\tau_0\tau_1 & \tau_1^2 \end{pmatrix} \right\}.$$

We also define state $l$, such that we use $\pi_{lj}$ to represent $1 - \pi_{\text{off},j} - \pi_{\text{on},j}$, $\pi_{\text{off},j}$, and $\pi_{\text{on},j}$ respectively for $l = 0, 1, 2$, for ease of exposition in these derivations. Similarly, $f_{lj}$ denotes the density associated with state

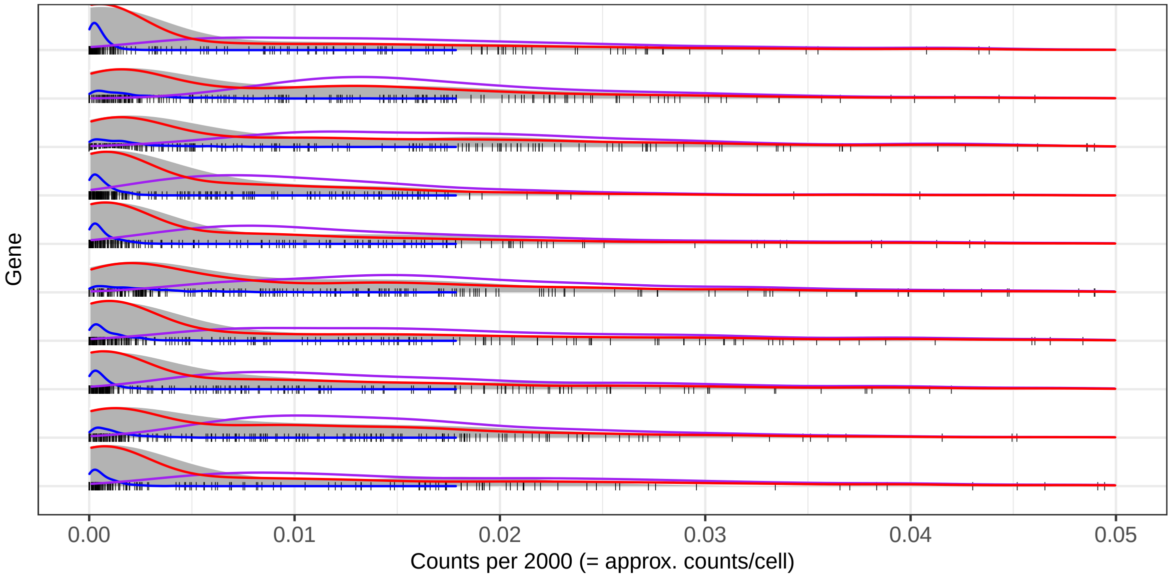


Figure 2: Fitted exponential and log-normal mixtures for genes determined to be unexpressed that demonstrate two off states. The blue densities represent the exponential components (off-low), and the purple densities represent the log-normal components (off-high). The red density represents the mixture of the two.

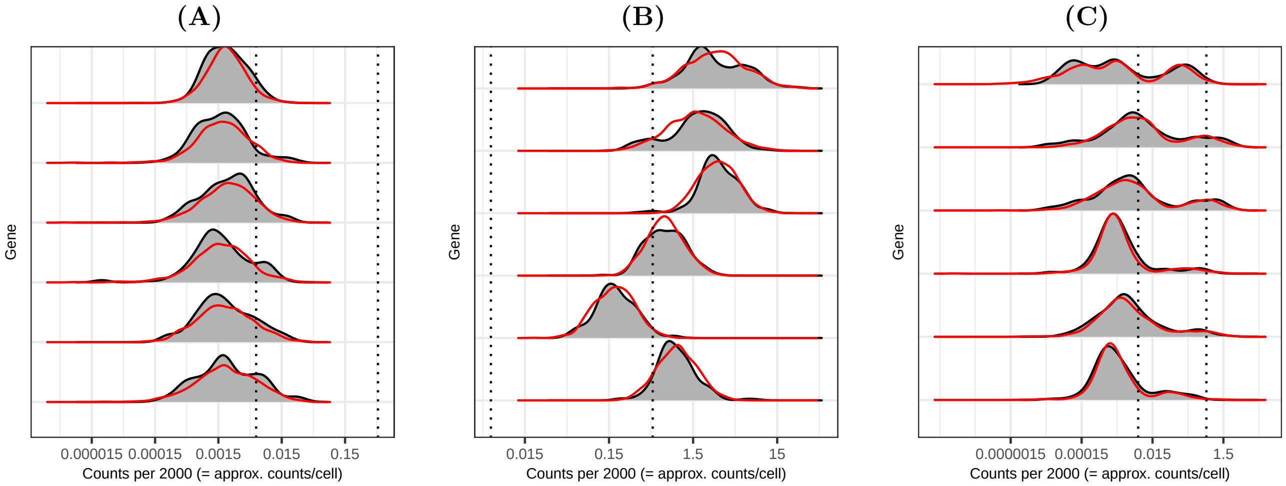


Figure 3: Fitted gene-specific distributions for examples of (A) genes that are estimated to be off in all cell- and tissue-types, (B) genes that are estimated to be on in all cell- and tissue-types, and (C) genes with estimated bimodal expression patterns. The red densities represent draws from the fitted mixture for each gene and are laid over the gray densities of the observed rates across cell- and tissue-types. The dotted lines represent the global off and on means respectively.

$l$ and gene $j$, and $Z_{jkl} = 1$ if gene $j$ belongs to state $l$ in cell-type $k$. Finally, we define a parameter vector $\boldsymbol{\theta}_j$ containing all the parameters we need to estimate for gene $j$, i.e. $\boldsymbol{\theta}_j = (\delta_{0j}, \delta_{1j}, \sigma_{0j}, \sigma_{1j}, \pi_{0j}, \pi_{1j}, \pi_{2j})$.

For the E-step, our goal is to compute the expected value of the complete-data log-posterior

$$Q(\boldsymbol{\theta}_j, \boldsymbol{\theta}_{j,curr}) = \mathbb{E}(\log \mathcal{L}(\boldsymbol{\theta}_j | \boldsymbol{\lambda}_j, \boldsymbol{Z}_j) | \boldsymbol{\lambda}_j, \boldsymbol{\theta}_{j,curr}) + \log \pi(\boldsymbol{\theta}_j).$$

Since this conditional expectation is over $\boldsymbol{Z}_j$, we can expand the first term as

$$\begin{aligned}\mathbb{E}(\log \mathcal{L}(\boldsymbol{\theta}_j|\boldsymbol{\lambda}_j, \boldsymbol{Z}_j)|\boldsymbol{\lambda}_j, \boldsymbol{\theta}_{j,curr}) &= \mathbb{E}\left(\sum_k \sum_l Z_{jkl} \log(\pi_{lj} f_{lj}(\lambda_{jk}|\boldsymbol{\theta}_j))|\boldsymbol{\lambda}_j, \boldsymbol{\theta}_{j,curr}\right) \\ &= \sum_k \sum_l \log(\pi_{lj} f_{lj}\left(\lambda_{jk}|\boldsymbol{\theta}_j)\right) \cdot \mathbb{E}(Z_{jkl}|\boldsymbol{\lambda}_j, \boldsymbol{\theta}_{j,curr}).\end{aligned}$$

Note that

$$\mathbb{E}(Z_{jkl}|\boldsymbol{\lambda}_j, \boldsymbol{\theta}_{j,curr}) = P(Z_{jkl} = 1|\boldsymbol{\lambda}_j, \boldsymbol{\theta}_{j,curr}) = P(Z_{jkl} = 1|\lambda_{jk}, \boldsymbol{\theta}_{j,curr}),$$

which we can compute using Bayes rule. In particular,

$$\begin{aligned}P(Z_{jkl} = 1|\lambda_{jk}, \boldsymbol{\theta}_{j,curr}) &= \frac{f(\lambda_{jk}|\boldsymbol{\theta}_{j,curr}, Z_{jkl} = 1)P(Z_{jkl} = 1|\boldsymbol{\theta}_{j,curr})}{f(\lambda_{jk}|\boldsymbol{\theta}_{j,curr}, Z_{jkl} = 1)P(Z_{jkl} = 1|\boldsymbol{\theta}_{j,curr}) + f(\lambda_{jk}|\boldsymbol{\theta}_{j,curr}, Z_{jkl} = 0)P(Z_{jkl} = 0|\boldsymbol{\theta}_{j,curr})} \\ &= \frac{\pi_{lj} f_{lj}(\lambda_{jk}|\boldsymbol{\theta}_{j,curr})}{\sum_{l'} \pi_{l'j} f_{l'j}(\lambda_{jk}|\boldsymbol{\theta}_{j,curr})}.\end{aligned}$$

We denote this as $\gamma(Z_{jkl})$. Since all other values in our expression are constant with respect to the expectation, we only need to compute $\gamma(Z_{jkl})$ in this E-step. We can then express

$$Q(\boldsymbol{\theta}_j, \boldsymbol{\theta}_{j,curr}) = \log \pi(\boldsymbol{\theta}_j) + \sum_k \sum_l \gamma(Z_{jkl}) \log(\pi_{lj} f_{lj}(\lambda_{jk}|\boldsymbol{\theta}_j)).$$

For the M-step, we now identify the values of $\boldsymbol{\theta}_j$ maximizing this expression.

For $\alpha_j$:

$$\begin{aligned}0 &\doteq \frac{\partial}{\partial \alpha_j} Q(\boldsymbol{\theta}_j, \boldsymbol{\theta}_{j,curr}) \\ &= \sum_k \gamma(Z_{jk0}) \frac{\partial}{\partial \alpha_j} \log\left(\pi_{0j} \cdot \frac{1}{\alpha_j} \exp\left(-\frac{\lambda_{jk}}{\alpha_j}\right)\right) \\ &= \sum_k \gamma(Z_{jk0}) \cdot \frac{-\frac{\pi_{0j}}{\alpha_j^2} \exp\left(-\frac{\lambda_j k}{\alpha_j}\right) + \frac{\pi_{0j}\lambda_{jk}}{\alpha_j^3} \exp\left(-\frac{\lambda_{jk}}{\alpha_j}\right)}{\frac{\pi_{0j}}{\alpha_j} \exp\left(-\frac{\lambda_{jk}}{\alpha_j}\right)} \\ &= \sum_k \gamma(Z_{jk0}) \left(\frac{1}{\alpha_j^2} - \frac{\lambda_{jk}}{\alpha_j}\right),\end{aligned}$$

which can be rearranged as

$$\sum_k \gamma(Z_{jk0})\lambda_{jk} = \frac{1}{\alpha_j} \sum_k \gamma(Z_{jk0}) \implies \alpha_{j,new} = \frac{\sum_k \gamma(Z_{jk0})}{\sum_k \gamma(Z_{jk0})\lambda_{jk}}.$$

For $\sigma_{0j}, \sigma_{1j}$:

$$
\begin{aligned}
0 &\doteq \frac{\partial}{\partial \sigma_{0j}} Q(\boldsymbol{\theta}_j, \boldsymbol{\theta}_{j,curr}) \\
&= \sum_k \gamma(Z_{jk1}) \frac{\partial}{\partial \sigma_{0j}} \log(\pi_{1j} f_{1j}(\lambda_{jk}|\boldsymbol{\theta}_j)) \\
&= \sum_k \gamma(Z_{jk1}) \frac{\partial}{\partial \sigma_{0j}} \log\left(\pi_{1j} \cdot \frac{1}{\lambda_{jk}\sigma_{0j}\sqrt{2\pi}} \exp\left(-\frac{(\log \lambda_{jk} - \mu_0 - \delta_{0j})^2}{2\sigma_{0j}^2}\right)\right) \\
&= \sum_k \gamma(Z_{jk1}) \cdot \frac{-\frac{\pi_{1j}}{\lambda_{jk}\sigma_{0j}^2\sqrt{2\pi}} \exp\left(-\frac{(\log \lambda_{jk} - \mu_0 - \delta_{0j})^2}{2\sigma_{0j}^2}\right) + \frac{\pi_{1j}}{\lambda_{jk}\sigma_{0j}\sqrt{2\pi}} \cdot \frac{(\log \lambda_{jk} - \mu_0 - \delta_{0j})^2}{\sigma_{0j}^3} \exp\left(-\frac{(\log \lambda_{jk} - \mu_0 - \delta_{0j})^2}{2\sigma_{0j}^2}\right)}{\pi_{1j} \cdot \frac{1}{\lambda_{jk}\sigma_{0j}\sqrt{2\pi}} \exp\left(-\frac{(\log \lambda_{jk} - \mu_0 - \delta_{0j})^2}{2\sigma_{0j}^2}\right)} \\
&= \sum_k \gamma(Z_{jk1}) \left(-\frac{1}{\sigma_{0j}} + \frac{(\log \lambda_{jk} - \mu_0 - \delta_{0j})^2}{\sigma_{0j}^3}\right),
\end{aligned}
$$

which can be rearranged as

$$
\sum_k \gamma(Z_{jk1}) = \frac{1}{\sigma_{0j}^2} \sum_k \gamma(Z_{jk1}) (\log \lambda_{jk} - \mu_0 - \delta_{0j})^2 \implies \sigma_{0j,new} = \sqrt{\frac{\sum_k \gamma(Z_{jk1})(\log \lambda_{jk} - \mu_0 - \delta_{0j})^2}{\sum_k \gamma(Z_{jk1})}}.
$$

Due to our constraints, we specifically update as

$$
\sigma_{0j,new} = \max\left\{0.5, \sqrt{\frac{\sum_k \gamma(Z_{jk1})(\log \lambda_{jk} - \mu_0 - \delta_{0j})^2}{\sum_k \gamma(Z_{jk1})}}\right\}.
$$

Symmetrically, we update

$$
\sigma_{1j,new} = \max\left\{0.5, \sqrt{\frac{\sum_k \gamma(Z_{jk2})(\log \lambda_{jk} - \mu_1 - \delta_{1j})^2}{\sum_k \gamma(Z_{jk2})}}\right\}.
$$

For $\delta_{0j}, \delta_{1j}$:

$$
\begin{aligned}
0 &\doteq \frac{\partial}{\partial \delta_{0j}} Q(\boldsymbol{\theta}_j, \boldsymbol{\theta}_{j,curr}) \\
&= \frac{\partial}{\partial \delta_{0j}} \log \pi(\boldsymbol{\theta}_j) + \sum_k \gamma(Z_{jk1}) \frac{\partial}{\partial \delta_{0j}} \log(\pi_{1j} f_{1j}(\lambda_{jk}|\boldsymbol{\theta}_j)) \\
&= -\frac{\partial}{\partial \delta_{0j}} \left(\frac{1}{2(1-\rho^2)} \left(\frac{\delta_{0j}^2}{\tau_0^2} - 2\rho \frac{\delta_{0j}\delta_{1j}}{\tau_0\tau_1} + \frac{\delta_{1j}^2}{\tau_1^2}\right)\right) + \\
&\qquad \sum_k \gamma(Z_{jk1}) \frac{\partial}{\partial \delta_{0j}} \log\left(\pi_{1j} \cdot \frac{1}{\lambda_{jk}\sigma_{0j}\sqrt{2\pi}} \exp\left(-\frac{(\log \lambda_{jk} - \mu_0 - \delta_{0j})^2}{2\sigma_{0j}^2}\right)\right) \\
&= -\frac{2\delta_{0j}}{2(1-\rho^2)\tau_0^2} + \frac{2\rho\delta_{1j}}{2(1-\rho^2)\tau_0\tau_1} + \sum_k \gamma(Z_{jk1}) \cdot \frac{\pi_{1j} \cdot \frac{1}{\lambda_{jk}\sigma_{0j}\sqrt{2\pi}} \cdot \frac{\log \lambda_{jk} - \mu_0 - \delta_{0j}}{\sigma_{0j}^2} \exp\left(-\frac{(\log \lambda_{jk} - \mu_0 - \delta_{0j})^2}{2\sigma_{0j}^2}\right)}{\pi_{1j} \cdot \frac{1}{\lambda_{jk}\sigma_{0j}\sqrt{2\pi}} \exp\left(-\frac{(\log \lambda_{jk} - \mu_0 - \delta_{0j})^2}{2\sigma_{0j}^2}\right)} \\
&= -\frac{\delta_{0j}}{(1-\rho^2)\tau_0^2} + \frac{\rho\delta_{1j}}{(1-\rho^2)\tau_0\tau_1} + \sum_k \gamma(Z_{jk1}) \cdot \frac{\log \lambda_{jk} - \mu_0 - \delta_{0j}}{\sigma_{0j}^2} \\
&= -\delta_{0j} \left(\frac{1}{(1-\rho^2)\tau_0^2} + \sum_k \frac{\gamma(Z_{jk1})}{\sigma_{0j}^2}\right) + \frac{\rho\delta_{1j}}{(1-\rho^2)\tau_0\tau_1} + \sum_k \gamma(Z_{jk1}) \cdot \frac{\log \lambda_{jk} - \mu_0}{\sigma_{0j}^2},
\end{aligned}
$$

which can be rearranged as

$$\delta_{0j,new} = \frac{\frac{\rho\delta_{1j}}{(1-\rho^2)\tau_0\tau_1} + \sum_k \gamma(Z_{jk1}) \cdot \frac{\log\lambda_{jk} - \mu_0}{\sigma_{0j}^2}}{\frac{1}{(1-\rho^2)\tau_0^2} + \sum_k \frac{\gamma(Z_{jk1})}{\sigma_{0j}^2}}.$$

Due to our constraints, we specifically update as

$$\delta_{0j,new} = \min\left\{\mu_1 + \delta_{1j} - \mu_0, \frac{\frac{\rho\delta_{1j}}{(1-\rho^2)\tau_0\tau_1} + \sum_k \gamma(Z_{jk1}) \cdot \frac{\log\lambda_{jk} - \mu_0}{\sigma_{0j}^2}}{\frac{1}{(1-\rho^2)\tau_0^2} + \sum_k \frac{\gamma(Z_{jk1})}{\sigma_{0j}^2}}\right\}.$$

Symmetrically, we update

$$\delta_{1j,new} = \max\left\{\mu_0 + \delta_{0j} - \mu_1, \frac{\frac{\rho\delta_{0j}}{(1-\rho^2)\tau_0\tau_1} + \sum_k \gamma(Z_{jk2}) \cdot \frac{\log\lambda_{jk} - \mu_1}{\sigma_{1j}^2}}{\frac{1}{(1-\rho^2)\tau_1^2} + \sum_k \frac{\gamma(Z_{jk2})}{\sigma_{1j}^2}}\right\}.$$

For $\pi_{0j}, \pi_{1j}, \pi_{2j}$:
We can use Lagrange multipliers as follows:

$$\begin{aligned}
0 &\doteq \frac{\partial}{\partial\pi_{lj}}\left(Q(\boldsymbol{\theta}_j, \boldsymbol{\theta}_{j,curr}) + \lambda\left(\sum_{l'=1}^{3}\pi_{l'j} - 1\right)\right) \\
&= \lambda + \sum_k \gamma(Z_{jkl})\frac{\partial}{\partial\pi_{lj}}\log(\pi_{lj} f_{lj}(\lambda_{jk}|\boldsymbol{\theta}_j))) \\
&= \lambda + \sum_k \gamma(Z_{jkl}) \cdot \frac{f_{lj}(\lambda_{jk}|\boldsymbol{\theta}_j)}{\pi_{lj} f_{lj}(\lambda_{jk}|\boldsymbol{\theta}_j)} \\
&= \lambda + \sum_k \gamma(Z_{jkl}) \cdot \frac{1}{\pi_{lj}},
\end{aligned}$$

which can be rearranged as

$$0 = \pi_{lj}\lambda + \sum_k \gamma(Z_{jkl}) \implies 0 = \lambda\sum_l \pi_{lj} + \sum_l\sum_k \gamma(Z_{jkl}) \implies \lambda = -K.$$

If we plug this back in, we find

$$\pi_{lj,new} = \frac{\sum_k \gamma(Z_{jkl})}{K}.$$
